# Neurobiological Basis of Brain Blood Oxygenation Responses Correlated with Cognitive Stroop Task Performance Before and After an Acute Bout of Aerobic Exercise

**DOI:** 10.1101/359307

**Authors:** Amrita Pal

**Author notes:** Corresponding author. E-mail address (A. Pal). ^1^. **Abbreviations** PFC: Prefrontal cortex MC: Motor cortex ER: Error Rate in Stroop Task RT: Reaction Time to Stroop Task HR: Heart Rate Hb: Hemoglobin Oxy-Hb (oxygen delivery, systemic) Deoxy-Hb (oxygen extraction, neuronal and glial) Total-Hb (cerebral blood volume, systemic) Diff-Hb (cortical oxygenation, neuronal).

## Abstract

Cardiovascular activities may increase the brain blood flow improving neural activities leading to improved cognition. Consequently, the effects of an acute bout of moderate intensity aerobic exercise on cortical brain blood oxygenation and its correlation with cognitive color-word Stroop task performance were tested. The Stroop tasks were congruent (color matches word) and incongruent (color does not match word). Prefrontal (PFC) and motor cortex (MC) blood flow was recorded by fNIRS (functional Near-Infrared Spectroscopy) while the human subjects performed the Stroop tasks before and after 30 minutes of exercise or equivalent time of rest (controlling for practice effect of repeating the Stroop task multiple times). It was predicted that PFC blood flow increase after exercise will correlate with increased Stroop interference and decreased cognitive flexibility after an acute bout of aerobic exercise at 70% of maximal Heart Rate (HR). We observed that Stroop interference in aerobic exercise and practice alone (rest) were not significantly (p>0.6) different indicating that exercise did not improve cognition above the learning effect. Increase in blood flow (both neurovascular and neurometabolic coupling) in previously deactivated PFC channels was positively correlated (p<0.05) to increased Stroop interference (worse cognitive speed and accuracy) after an acute bout of aerobic exercise.

**Highlights:**

- Does moderate intensity aerobic exercise improve cognition?
- Does prefrontal cortex oxygenation correlate with cognition changes after exercise?

## 1. Introduction

### Aerobic exercise and cognition

Studies confirmed that moderate intensity aerobic exercise improved response-inhibition involving cognitive task speed by 22%, but accuracy was not affected (Rattray et al., 2013). In soccer players, increased aerobic exercise intensity did not affect accuracy of visual-attention decision making tasks, but did improve the motor RTs by 2.29% (Frýbort et al., 2016). Increased submaximal aerobic exercise intensities led to exercise induced physiological arousal (sweating, higher heartbeat, etc.) in soccer players (experienced or inexperienced), which led to a 14% improvement in the speed of decision making but no change in accuracy (Fontana et al., 2009). Higher plasma norepinephrine and adrenocorticotropic hormone concentrations correlate with a decline in executive function after high intensity exercise (75% of maximal exercise intensity; Konishi et al., 2017). Overall, moderate but not high intensity exercise appears to facilitate cognitive speed, but does not improve accuracy of decision making.

To establish if increase in blood flow after exercise improves cognition, it is important to first find correlations between changes in HR and behavioral data before we could correlate brain blood flow to behavioral data. Important to note though that changes in HR is not equivalent to changes in the cardiac output (cardiac output = Heart Rate X Stroke Volume) (Lee & Oh, 2016). Nor are changes in cardiac output directly correlated with changes in the neural activity, although there were evidences of neurovascular coupling (Woolsey et al., 1996). Conflict-resolution aspect of decision making as measured by Stroop tasks.

The color-word Stroop task (Stroop, 1935) tests for selective attention to specific information during decision making and involves inhibition of conflicting responses (interferences) (MacLeod, 1991). An example of incongruence in the color-word Stroop task has the subject read the color and not the semantics of the word presented. Interference occurs when there is competition between conflicting choices of color and word. The Stroop effect is a highly reproducible effect and is associated with the longer RT in the incongruent task compared to the congruent task (color and word match) (MacLeod, 1991). The brain may use a mean firing rate of a group of neurons with similar functions for the decision-making task, or the brain may use a linear combination of firing rates of many different neurons and their interactions or projections to one another (output) for making the final decision (Beck et al, 2008). In this project, we predict that the encoding schema used is scalar population rate coding (Van Opstal, 2007; Tranquillo, 2008).

### functional Near Infrared Spectroscopy (fNIRS)

The light frequencies at infrared and near-infrared wavelengths are used to differentiate the relative levels of oxy-Hb and deoxy-Hb in the cortical tissue (Plenger et al., 2016; Tam & Zouridakis 2014; Yamada et al., 2012). In this technique, light is directed into the brain tissue by fibre-optic bundles called optodes (emitters). A second set of optodes (detectors) collect light after it has passed through the tissue (Tam & Zouridakis 2014; Villringer & Chance, 1997). Typically, the emitters and detectors are placed on wearable headcaps (Tam & Zouridakis 2014; Villringer & Chance, 1997). Increases in number of source-detector distances enabled higher depth sensitivity up to the level of detection in the sulcal folds (Scholkmann et al., 2014). The modified Beer-Lambert law is used to calculate the optical density or absorption changes based on light scattering through the brain tissue, after filtering the raw data for cardiac oscillations (Scholkmann et al., 2014; Schytz, et al., 2009; Butti et al. et al., 2007; Delpy et al., 1988). fNIRS gives insight into the metabolic rate of the cortical pyramidal neuronal and glial responses, through the brain blood oxygenation changes in the prefrontal or motor cortices (Plenger et al., 2016; Tam & Zouridakis 2014; Yamada et al., 2012; Merzagora et al., 2009). Brain activation using the fNIRS technique is deduced from an increase in oxy-Hb and a simultaneous decrease in deoxy-Hb during an excitatory cognitive task (Vazquez et al., 2010; Schecklmann et al., 2008).

fNIRS technology has been widely used for quantifying Stroop task response in the brain (Plenger et al., 2016; Lambrick et al., 2016; Tam & Zouridakis 2014; Byun et al., 2014). Usually non-parametric correlations are used to correlate the non-normally distributed Stroop task performance data with normally distributed brain blood oxygenation data (Yanagisawa et al., 2010). Oxy-Hb increased during the incongruent Stroop task compared to the congruent Stroop task, measured bilaterally at the frontal lobes (dorsolateral PFC) (Plenger, 2016). In addition, higher oxy-Hb at the left somatosensory and motor cortices correlated with the act of vocalizing the responses during the Stroop tasks (Plenger et al., 2016).

Acute exercise bouts with progressively higher exercise intensities (52%, 68%, 84%) increased both oxy and deoxy-Hb (overall oxygenation) at the PFC (Giles et al., 2014). In children aged 8-10 years involved in moderate intensity aerobic activity, all three variables of oxy, deoxy and total-Hb showed increases, yet, total-Hb (total perfusion) accounted for the highest variance (38%) post-exercise corresponding to improvement in Stroop performance (Lambrick et al., 2016). Oxy-Hb increase was attributed to the apparent local rise in cerebral blood flow in excess of oxygen demand and higher cardiac output post submaximal exercise, contributing to improved cognition (Ide et al., 2000). If neuronal or glial activity increases after exercise, oxygen extraction will increase (Vazquez et al., 2010). Thus, the deoxy-Hb (oxygen extraction, neuronal or glial) should go higher even if the oxy-Hb (oxygen delivery, systemic) may not keep up with this oxygen demand, given that limited metabolic resources are also going to the high-performing muscle tissues during exercise (Tam & Zourdakis, 2014). However, what we observed in Lucas et al. (2012), Byun et al. (2014) and Yanagisawa et al. (2010) the deoxy-Hb dropped with higher exercise intensities. Initially, at 30% aerobic exercise intensity, subjects made more errors (maybe a sympathetic response), however, at 70% exercise intensity, subjects made fewer errors compared to resting baseline, both for the congruent and incongruent Stroop tasks (Lucas at al., 2012). The deoxy-Hb at the right PFC increased from the resting levels when exercised at 30% of maximal heart rate, however, at 70% of maximal heart rate, deoxy-Hb levels returned towards, but did not fully reach resting baselines (Lucas et al., 2012). Oxy-Hb had the opposite response to deoxy-Hb, in that, there was an initial decrease for up to 30% of exercise intensity and then a gradual increase for up to 70% at the PFC (Lucas et al., 2012). After an acute bout of 10 minutes of mild aerobic exercise at 30% maximal heart rate, fNIRS revealed higher cortical activations in terms of higher oxy-Hb (with no significant deoxy-Hb changes) evoked in the left dorsolateral (DLPFC) and left frontopolar regions (Byun et al., 2014). With an acute bout of exercise at 50% of maximal heart rate, subjects showed an increase in oxy-Hb but not deoxy-Hb at the left DLPFC (Yanagisawa et al., 2010). The initial effects of moderate intensity exercise caused the uncoupling of the cortical blood oxygenation from the cognition. This effect was moderated only if there was enough oxygen delivery during the course of exercise with intensities as high as 80%. Moreover, if exercise became too difficult or at a supra-maximal intensity (>80%), then cognitive performance in Stroop tasks again declined (Ogoh et al., 2014). Thus, the alternate hypothesis was other mechanisms like release of neurotransmitters and not increased blood perfusion after exercise contributed to improved cognition (Meeusen & Meirleir, 1995).

### Predictive Model

We predicted that after an acute bout of sub-maximal aerobic exercise compared to practice alone (rest), there would be no Stroop interference improvement, and this would correlate with increase in cortical oxygenation in prefrontal cortex regions that show decreased oxygenation (neural activity) before exercise.

## 2. Material and methods

### Participants

The Institutional Review Board (IRB) guidelines of University of North Texas UNT Denton, (IRB #14453) were followed in the recruitment of the subjects. Adults signed an informed consent form and, for minors, their guardian signed a consent form. Refer to Table 1 for the demographic distribution of the subjects. We had to eliminate 1 subject from Rest group and 3 subjects from Exercise group since fNIRS data collection was poor as evidenced by extremely large effect size (>50) changes in task signals from baseline.

**Table 1.** Participants.

| Paradigm | n | Age | Gender |
| --- | --- | --- | --- |
| Exercise | 73 | 23± 5 | 22 Males<br>51 Females |
| Non exercise | 30 | 23± 5 | 11 Males<br>18 Females |
| Overlap | 13 | 24± 6 | 4 Males<br>9 Females |

### Experimental Design

#### Rest (control)

As a control for the practice effect that would likely result from the Stroop tasks being conducted multiple times, subjects participated in separate, rest (control) in which they recorded the Stroop tasks before and after 30 min of rest without doing the experiment of exercise (Figure 1). If the same subjects who did the exercise experiments volunteered to be a part of rest (control), they returned on a later day. During the Stroop tasks, brain blood oxygenation was measured using fNIRS from the PFC and MC.

**Figure 1.**
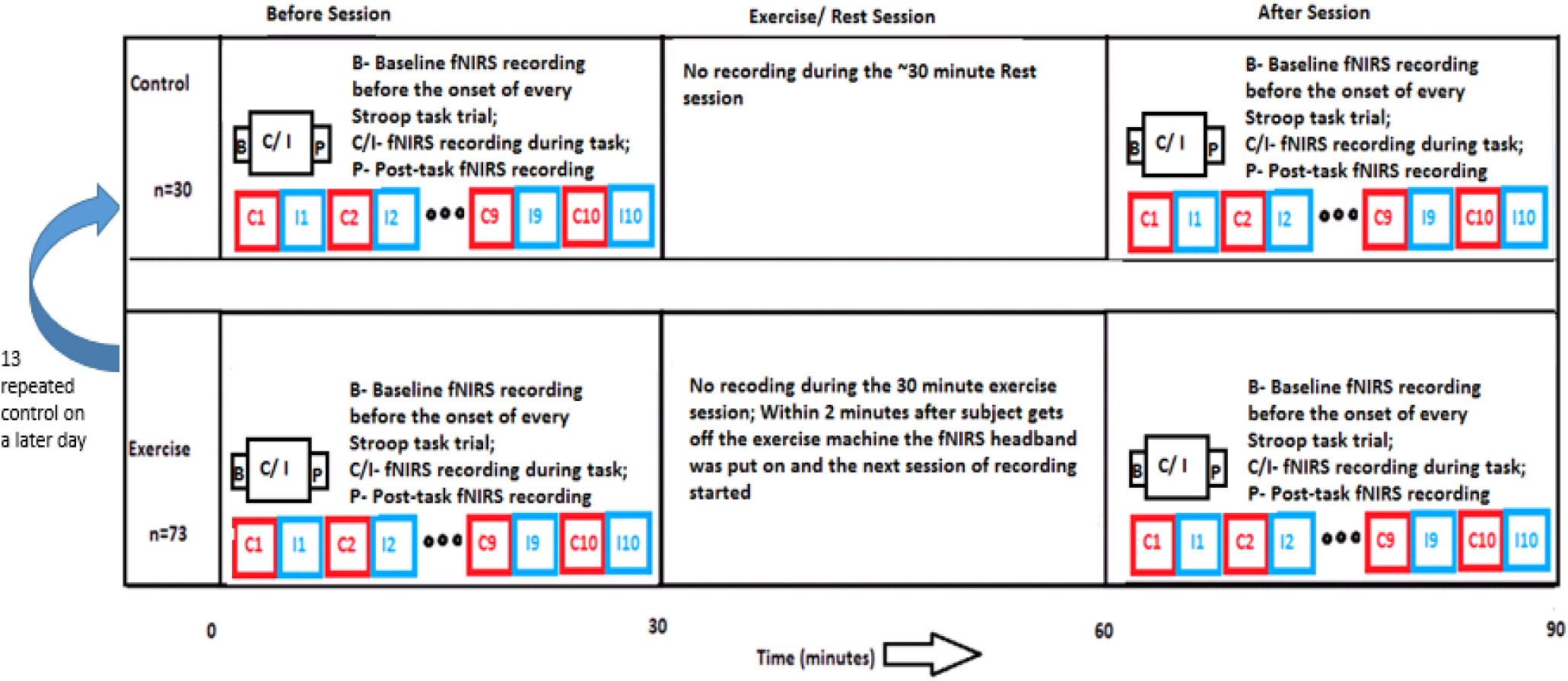
Overall design of the experiment. Each trial recording consisted of right before (baseline, B), during (task, C/I) and right after (post-task, P) the Stroop task. Twenty such recordings were made consecutively with congruent and incongruent trials alternating in a pseudo-randomized style in each session (before/after). During the intermittent rest or exercise session, no fNIRS recording was performed.

#### Exercise

Subjects performed the Stroop tasks before and after 30 minutes of aerobic exercise at moderate intensity (70% of maximal HR) performed on a stationary bike (Cardio Dual Trainer, BRM 3671/3681/3690) (Figure 1). 70% of the maximal heart rate was determined from the participants’ age, by subtracting the person’s age from 220 (Mikus et al., 2009). During the Stroop tasks, brain blood oxygenation was measured using fNIRS from the PFC and MC.

#### Stroop tasks

Subjects performed both the congruent and incongruent Stroop color-word tasks in pseudo-randomized alternating blocks consisting of 10 trials in each block (Figure 1). The subjects read aloud the color of 40 randomized printed words on a computer screen as fast as possible. The time for the completion of the above task was recorded by the webpage (http://biology.unt.edu/~tam/Science/StroopTest.html) when the start and end buttons were pressed at the beginning and end of task. The experimenter recorded the accuracy of the color-identification.

#### Time markers for fNIRS recordings during Stroop tasks

Time markers were recorded by the experimenter to synchronize with the onset and the end of the Stroop task for subsequent analyses (Figure 1). The neural recording consisted of a minimum of 5 sec and a maximum of 20 sec of resting level baseline, during which the subject was instructed to sit stationary and not engage in any mental or physical task. This was followed by administration of the Stroop task, typically ~40 secs for the incongruent Stroop task and ~20 secs for the congruent Stroop task. This was followed by another 5 sec to 20 sec of resting-level post-task recording after the Stroop task.

#### fNIRS equipment

Imagent fNIRS equipment used for the current study was a continuous wave system that detects statistically significant relative changes in brain oxy- and deoxy-Hb levels rather than quantifying it absolutely (Scholkmann et al., 2014). In this equipment, laser-diodes were light emitters and the detectors collected the light passed through the prefrontal and motor cortices. fNIRS data collection from the Boxy software associated with this device found the oxy- and deoxy-Hb levels from the light passed through these brain regions using the modified Beer-Lambert law. This software Boxy was calibrated each time a subject donned the fNIRS headcap immediately before the recording.

#### Optode Placement in fNIRS headcap

The Imagent manual (2005) provides the details of fNIRS as applied to the assessment of human brain. The fNIRS headcap was fastened to the subject’s head with Velcro straps, using the 10-20 International convention (Xiao et al, 2017) (Figure 2 A.). In total, 4 detectors and 24 channels were employed. The optode placements were shown in Figure 2 B. and C. The emitters and detectors were 2-4 cm apart. The optic channels from the emitters to the first set of detectors A and B were placed on the subject’s “Nasion Area”, in which, the left and right PFC were odd and even numbered channels respectively (Figure 2 B.). The second set of detectors C and D were placed on the subject’s “Left Pre-Auricular Area” of the left hemisphere if the subject was right-dominant, and vice-versa if he/she was left-dominant. The dorsal channels (C and D odd-numbered) monitored the pre-motor area and the ventral channels (C and D even-numbered) monitored the motor area. We evaluated the PFC channel activity per our predictive model.

**Figure 2.**
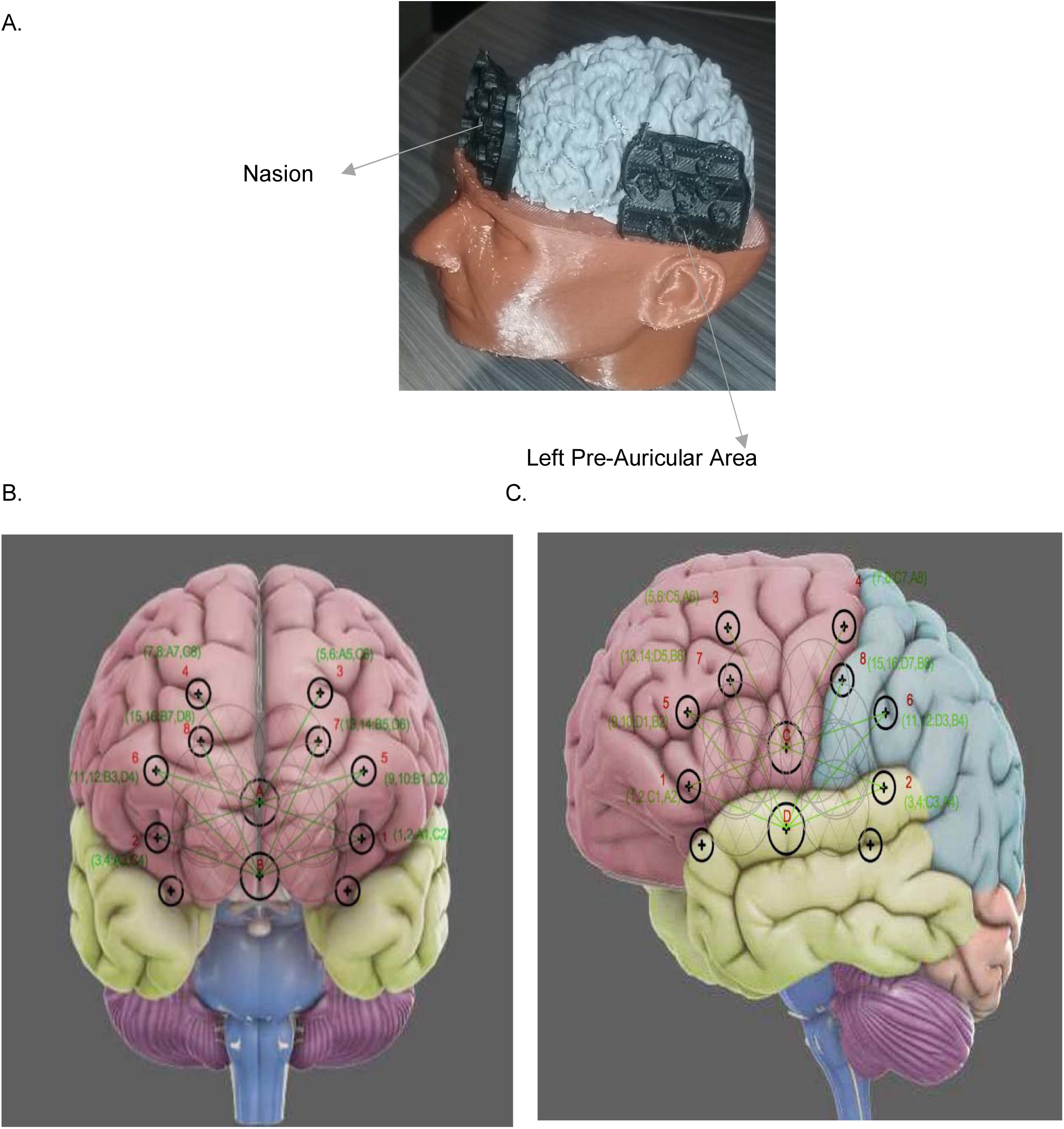
A. Placement of fNIRS cap at the Nasion to record PFC and Left Pre-Auricular Area to record MC, picture of 3D print created by Amrita Pal (2017). Temporal montage of the optode placement on the fNIRS Imagent cap on PFC (B.) and MC (C.), modified by Tam & Zouridakis (2014).

#### fNIRS Protocol

All subjects were instructed to minimize head movements. Lights in the recording room were turned off to prevent ambient light noise in the recordings. The Stroop task stimuli was at the center of the computer screen and easily visible to the subject, so that the optical recordings were not compromised by the Stroop task measurements. If a subject had thick black hair, which would interfere with the absorption of the infrared light, recordings were made from only the prefrontal cortex (hair free forehead) but not from the motor cortex. Hair was placed aside underneath the optical fibers whenever possible and comfortable. While the channel locations could be different with respect to the brain location from subject to subject, the relative changes after exercise or after rest for the general brain location did not vary from subject-to-subject. Thus, the physiological interpretations of the data were consistent across all subjects.

#### Statistics

Error rate in each trial was calculated as percentage of errors out of the possible 40 words per trial. The relative changes in error rate (ER) and reaction time (RT) in the 10 trials after exercise (or rest) compared to 10 trials before exercise (or rest) were obtained and the Stroop interference was calculated as the value for incongruent minus congruent. The relative changes in Stroop interference for the ER and RT was compared between the exercise and rest groups using independent samples t-test with unequal variance.

Also, change in heart rate (HR) after exercise (or rest) during incongruent minus congruent conditions were obtained as well. These relative changes (parametric) in HR were correlated with RT and ER using Pearson’s R correlation for the exercise and rest groups. The trends were compared for the exercise versus rest conditions.

ISS Imagent fNIRS equipment generated time series of oxy-Hb and deoxy-Hb data processed by the software Boxy. fNIRS signal variables (oxy-Hb, deoxy-Hb and derived measures of total-Hb, diff-Hb) for the 12 PFC channels were then estimated by a time-average (± SEM) for before the Stroop tasks began (Baseline), during the Stroop tasks (Task), and after the Stroop task ends (Post-task) in order to remove noise in the time series signals in each trial. These analyses were automated using VBA EXCEL Macro programming in all the subjects. The brain blood oxygenation data was then baseline corrected i.e. task and post-task time averages were deducted from the baseline time average, and then averaged over 10 trials in one subject for a specific experimental condition (congruent or incongruent, before or after, exercise or rest). To determine Task or (Post-task) responsive consistent change in fNIRS signal compared to Baseline, we calculated the effect size Cohen’s D (Lakens, 2013; Fritz et al., 2012; Cohen, 1988) for the 10 trials in each before experimental (exercise or rest) condition in each subject by calculating the average of the difference between Task (or Post-Task) minus Baseline for all 10 trials divided by the standard deviation pooled of all 10 Baseline in the 10 trials. We then determined the PFC channel (out of the possible 12) showing most consistent negative (and positive) task-responsive changes in Cohen’s D of incongruent minus congruent trials in each subject in the before exercise (or rest) conditions. Then for the same channels in the same subject that were most task responsive before exercise (or rest), we calculated the relative changes in the trial averaged fNIRS signals after exercise (or rest). These relative changes in the trial averaged fNIRS signals were then correlated with the relative changes in the RT and ER using Pearson’s R correlation in all subjects. We then compared the trends of the exercise experiment and the practice (rest) controls.

## 3. Results

### Behavioral Data

The Stroop interference change in RT (p=0.73) and ER (p=0.42) after exercise did not show any significant difference from rest (practice) controls (Figure 3).

**Figure 3.**
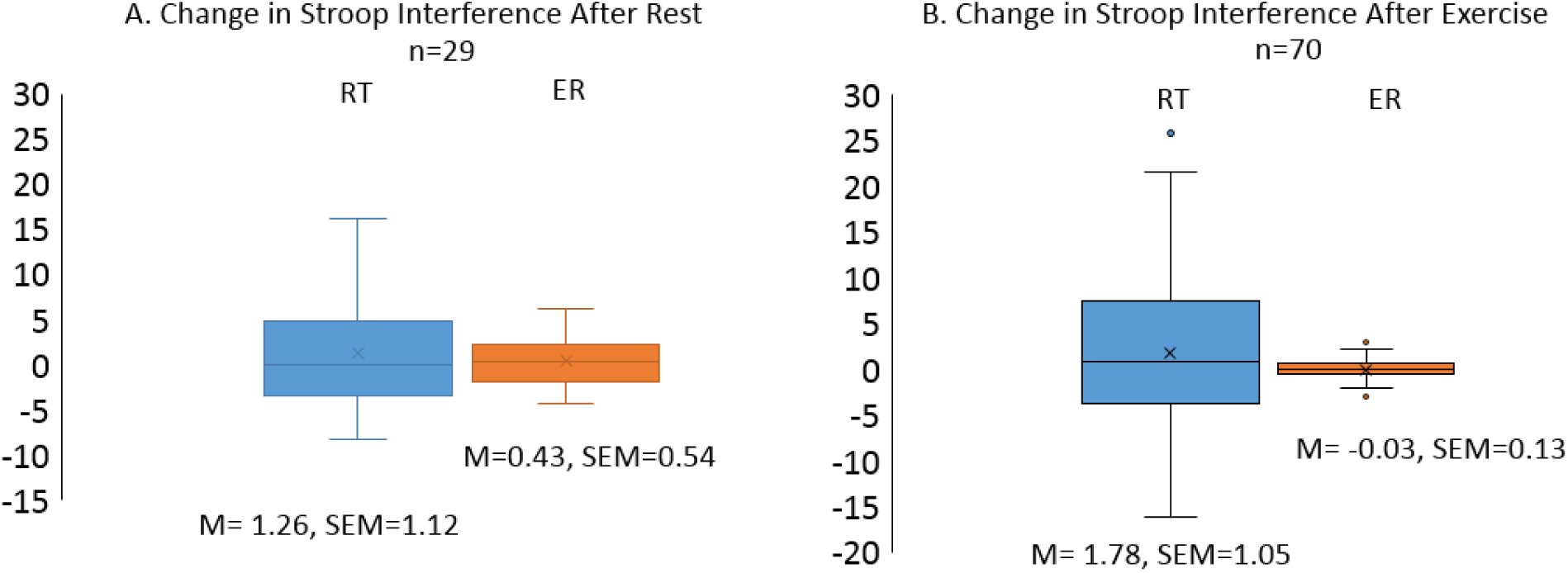
Percentage change in Stroop interference (incongruent minus congruent) of RT and ER after rest (A) and after exercise (B). There was no significant difference between Rest (practice or learning effect) and Exercise Stroop interference, p=0.73 for RT, p=0.42 for ER.

### HR Data

HR correlated with the ER for the rest group (p<0.1) but there was no significant correlation between HR and RT (p=0.79) for the rest group, also no correlation between HR and RT (p=0.86) and between HR and ER (p=0.86) in the experimental group (Figure 4).

**Figure 4.**
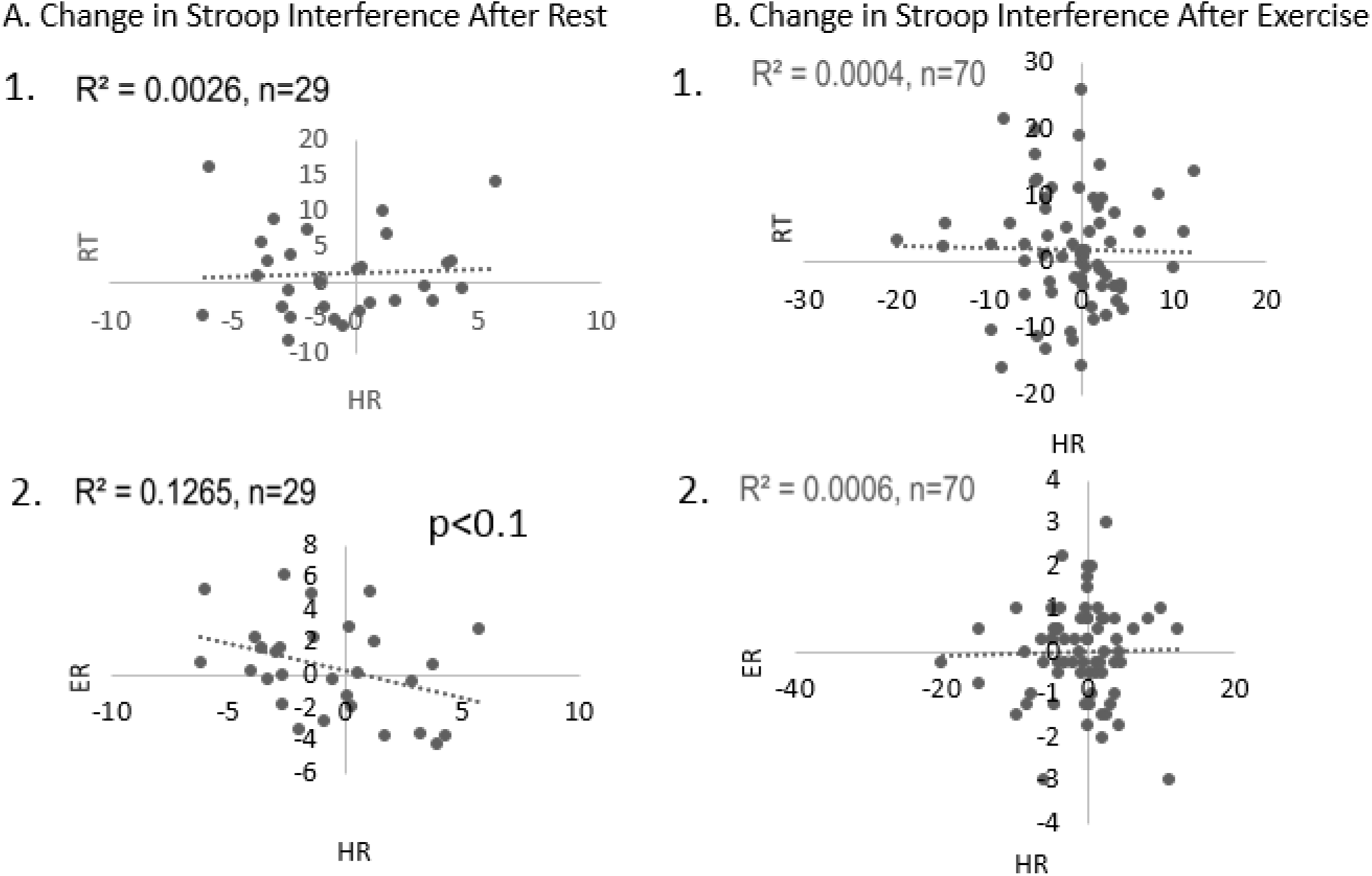
Correlation of change in Stroop interference (RT and ER) vs HR after rest (A) and exercise (B). Only ER correlate with HR after rest (p<0.1). No correlation between after exercise change in HR with RT or ER during incongruent-congruent trials.

### fNIRS Data

Distribution of the Cohen’s D values of the most task responsive PFC channels showing positive and negative changes in fNIRS signal from baseline in the incongruent minus congruent trials for the before rest and before exercise conditions (Figure 5). High effect size changes were mostly found in the oxy-Hb (neurovascular coupling) and then in the diff-Hb (neurometabolic coupling). For these specific channels, the trial-averaged fNIRS signal changes in incongruent minus congruent trials after exercise (or rest) were correlated with the changes in RT and ER in all subjects as shown in Figure 6. For PFC channels deactivated (negative Cohen’s D) before practice (rest), increase in oxygen delivery (systemic or neurovascular coupling) was positively correlated (p<0.05) to increase in Stroop interference in Error Rate (more errors) (Figure 6 A). For PFC channels initially activated (positive Cohen’s D) before rest, decrease in cortical oxygenation (neural activity or neurometabolic coupling) was positively correlated (p<0.05) to Stroop interference in Error Rate (more errors) but negatively correlated to Reaction Time (faster) indicating speed-accuracy tradeoff (Figure 6 A). Increase in prefrontal cortical oxygenation (neurometabolic and neurovascular coupling, increased blood flow) in PFC channels deactivated before exercise correlated positively (p<0.05) with increased Stroop interference (worse cognitive performance speed RT and accuracy ER) after exercise (Figure 6 B). For PFC channels initially activated before exercise there was no significant correlation (p>0.1) between change in oxygenation and behavioral data after exercise.

**Figure 5.**
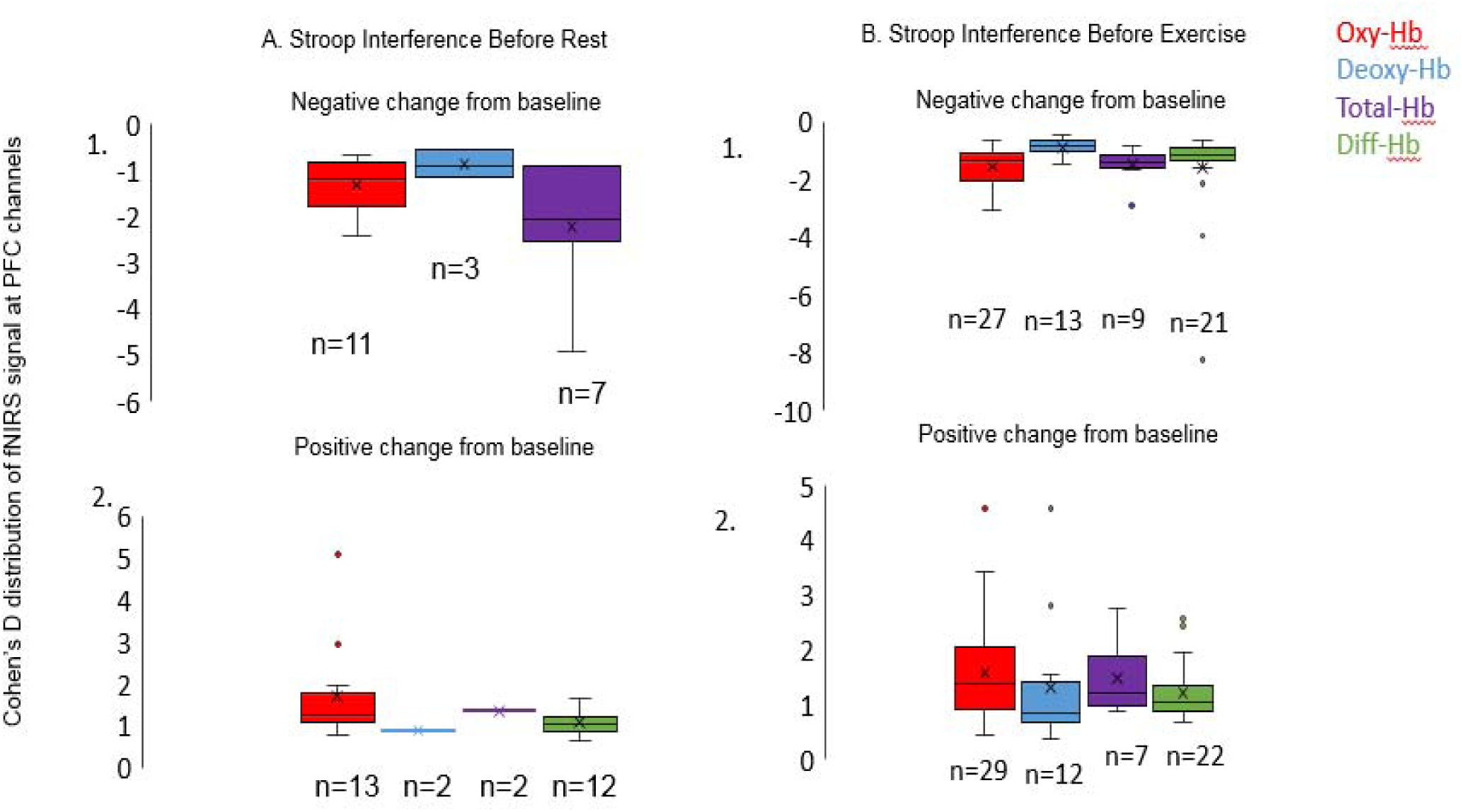
Negative and positive Cohen’s D distribution of fNIRS signal (red: oxy-Hb, blue: deoxy-Hb, purple: total-Hb and green: diff-Hb) from PFC channels showing Stroop interference (incongruent minus congruent) responses before rest (A) and before exercise (B) conditions.

**Figure 6.**
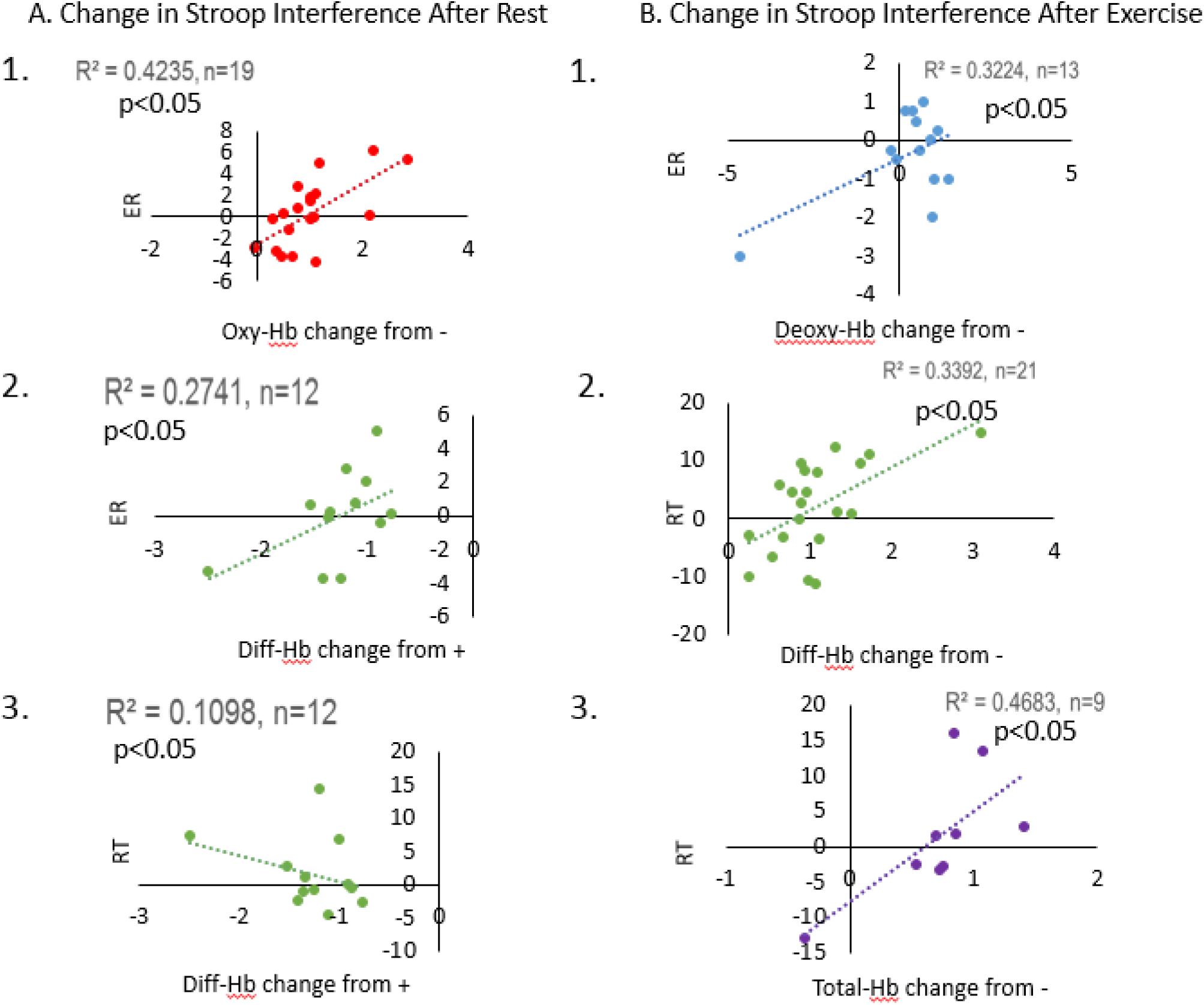
Summary of significant correlations (p<0.05) of change in fNIRS signals (incongruent minus congruent) with Stroop interference change in ER and change in RT after rest (A) and after exercise (B).

## 4. Discussion

After an acute bout of aerobic exercise there was no significant difference in Stroop effect compared to practice (learning effect) indicating report of Bergquist et al. (2013) and could be explained as an impact of sympathetic exercise induced arousal that decreased cognitive flexibility.

Change in HR during the incongruent minus congruent trials did not correlate with Stroop Interference measures of speed and accuracy after exercise (Figure 4). However, after rest, error correlated negatively with the HR indicating fewer errors when the HR goes up after rest (Figure 4).

We confirmed that the brain regions in the prefrontal cortex were participating (via activation or deactivation) in response to the Stroop tasks similar to what have been found earlier (Plenger et al., 2016; Giles et al., 2014). Majority of the subjects showed high effect size most-task responsive changes in the oxy-Hb signal and then diff-Hb signal (Figure 5). Thus, oxy-Hb changed rapidly rather than deoxy-Hb. This is explained as a metabolic lag where systemic changes (oxy-Hb) occurred first and had higher effect sizes changes from the baseline than the neural (deoxy-Hb) changes, which occurred later. The switch from mitochondrial to glycolytic machinery for neural firing (Jang et al., 2016) could explain the lag in the metabolic response for the deoxy-Hb compared to the oxy-Hb. When enough oxygen is being delivered to the brain tissue (oxy-Hb), the neurons utilized mitochondrial oxidative phosphorylation. Soon after, the energy mode shifted to being glycolytic as oxygen delivery was not sufficient for pertinent functional processing in the brain. The oxygen extraction variable (deoxy-Hb) variable then started changing. It also explains why previous studies reported only systemic (oxy-Hb) changes after exercise (Giles et al., 2014; Tempest et al., 2014; Lambrick et al., 2016; Byun et al., 2014; Yanagisawa et al., 2010) in response to the Stroop tasks (Plenger et al., 2016). To confirm if the changes were truly systemic or neural or both, the later tests on correlation of performance in terms of accuracy and RT to the brain blood oxygenation was conducted for each of the exercise and rest groups. Increase in oxygen delivery (oxy-Hb) in previously deactivated PFC channels correlated positively to errors made after rest (Figure 6 A. 1.) indicating more errors were made when oxygen delivery increased in those channels. In addition, after rest, decrease in PFC oxygenation in previously activated channels correlated positively with errors made and correlated negatively with reaction time, indicating fewer errors and slower speed (speed-accuracy tradeoff) when cortical oxygenation decreased in previously activated channels (Figures 6 A. 2 and 3.). Increase in blood flow (both neurovascular and neurometabolic coupling) in previously deactivated PFC channels was positively correlated to increased Stroop interference (worse cognitive speed and accuracy) after an acute bout of aerobic exercise (Figure 6 B). Again, this could mechanistically explain why cognitive flexibility was compromised after an acute bout of aerobic exercise—the PFC channels possibly involved in impulsivity were turned on with the sympathetic arousal induced by exercise. Further research into the demographics (ADHD (Attention-Deficit Hyperactivity Disorder), obesity etc.) of the population demonstrating this response could explain if this negative impact on cognitive flexibility with aerobic exercise was especially detrimental to a particular sub-population.

## Acknowledgements

I want to acknowledge my mentor Dr. Nicoladie Tam who helped design the experiment and allowed me to record in her laboratory. I want to acknowledge my mentor Dr. Warren Burggren who helped me write the material, figures and proofread it for scientific and grammatical errors. Dr. Rich Herrington from UNT RSS (Research and Statistical Services) was very helpful in his contribution to the statistical data analysis. I thank Ala Obaid for her assistance in the data collection during some experiment sessions.

## Funding

This research did not receive any specific grant from funding agencies in the public, rcial, or not-for-profit sectors.

